# Detecting within-host interactions from genotype combination prevalence data

**DOI:** 10.1101/256586

**Authors:** Samuel Alizon, Carmen Lía Murall, Emma Saulnier, Mircea Sofonea

**Affiliations:** MIVEGEC, CNRS, IRD, Université de Montpellier, France

**Keywords:** multiple infections, MOI, superspreaders, inference, ABC, competition

## Abstract

Parasite genetic diversity can provide information on disease transmission dynamics but most methods ignore the exact combinations of genotypes in infections. We introduce and validate a new method that combines explicit epidemiological modelling of coinfections and regression Approximate Bayesian Computing (ABC) to detect within-host interactions. Using genital infections by different types of Human Papillomaviruses (HPVs) as a test case, we show that, if sufficiently strong, within-host parasite interactions can be detected from epidemiological data and that this detection is robust even in the face of host heterogeneity in behaviour. These results suggest that the combination of mathematical modelling and sophisticated inference techniques is promising to extract additional epidemiological information from existing datasets.

## Introduction

Hosts are known to often be simultaneously infected by multiple genotypes of the same parasite species or even by multiple parasite species. Over the last decades, the gap between our ability to detect this parasite within-host diversity and its use in epidemiological inference model has widened. Here, we introduce and validate an approach to detect within-host interaction from equilibrium prevalence data even in the presence of another source of heterogeneity. This method relies on the exact combination of parasite genotypes in each host, which we hereafter refer to as the ‘genotype combination’. We focus on genital infections by different types of human papillomaviruses (HPVs), which are known to be highly prevalent (Thomas et al., 2000, Rousseau et al., 2001, Chaturvedi et al., 2011), but this method is applicable to any system of multiple infection by different parasite species or genotypes for which there is sufficiently rich data.

### Binary or rank models

Most epidemiological models that allow for parasite genotypes to coexist within a host only allow for up to two genotypes per host and do not allow for cotransmission, although there are exceptions for both (May & Nowak, 1995, Lion, 2013, Alizon, 2013, Sofonea et al., 2015). These ‘binary’ models have been instrumental in epidemiology but are by definition inappropriate as soon as parasite diversity exceeds three genotypes.

Conversely, studies on macro-parasites have long been incorporating the multiplicity of infection in their models (Anderson & May, 1978). They showed that the distribution of the number of macro-parasites per host, which we here refer to as the ‘rank’ of an infection, can provide information regarding the contact structure within the host population. In absence of heterogeneity of any kind, one would expect rank distributions to follow a Poisson distribution. Interestingly, in many populations, the number of macro-parasites per host tends to follow a negative-binomial distribution, which is often interpreted as evidence for some sort of host population structure (Shaw & Dobson, 1995, Wilber et al., 2017). This aggregation pattern then shapes the functional response between parasitism and host death rate in ways that can critically affect population dynamics (Anderson & May, 1978).

For microparasites, similar studies have been developed, where the infection rank corresponds to the number of genotypes detected in a host. For example, Chaturvedi et al. (2011) showed that a Poisson distribution can be rejected for HPV genital infections suggesting that there is an excess of coinfections compared to what would be expected in a standard Susceptible-Infected (SI) model. Additional analyses of ours show that a negative binomial distribution nicely captures the tail of this distribution (Fig 1A). This is consistent with the fact that the ‘number of lifetime partners’ was the cofactor the most strongly associated with being infected by multiple HPV types instead of a single HPV type in the study by Chaturvedi et al.

**Fig 1.**
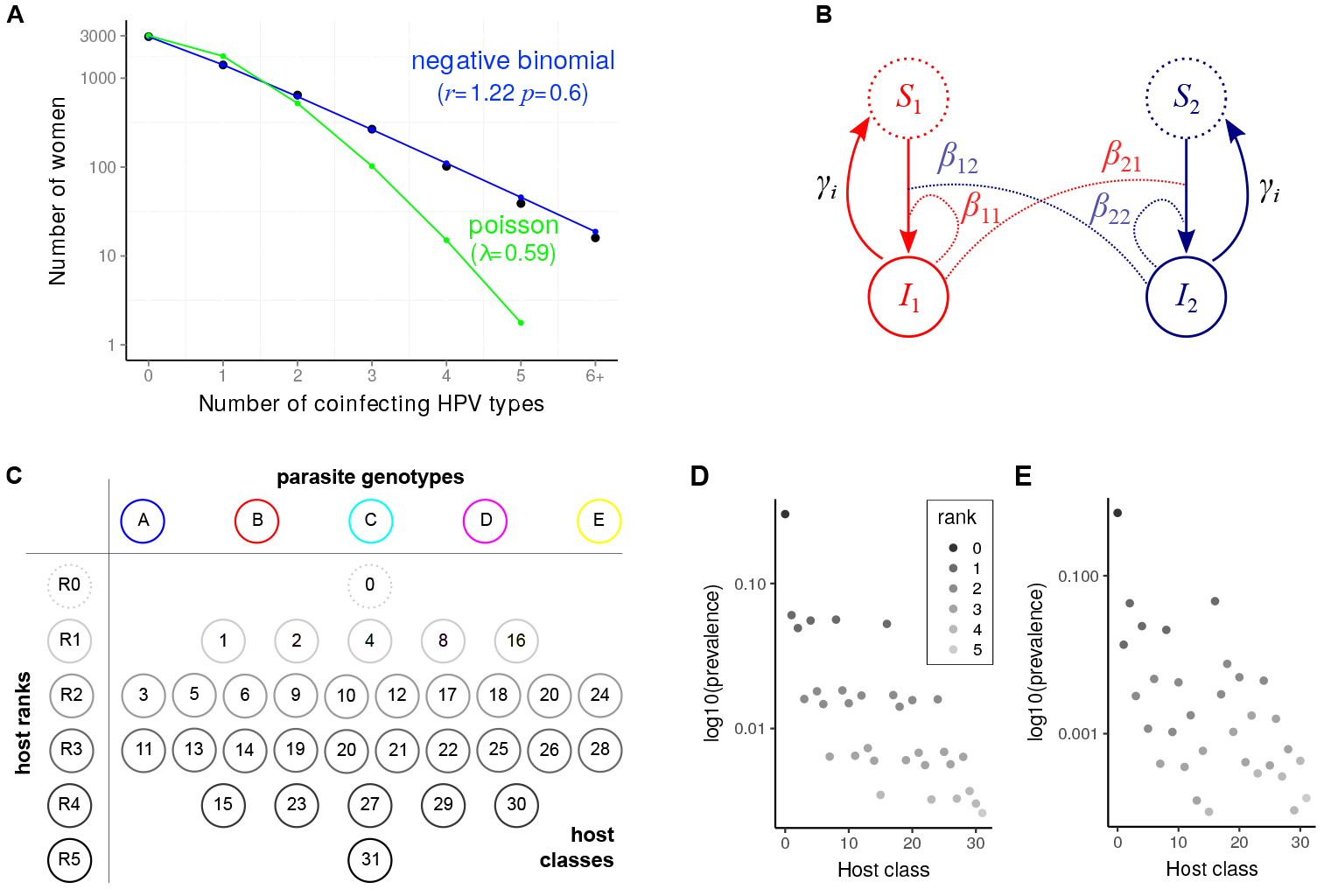
The coinfection epidemiological setting. A) Empirical rank distribution for HPV infections, B) Flow diagram showing the population structure with ‘normal-spreader’ (1 in red) and ‘super-spreader’ hosts (2 in dark blue), C) Host class prevalences for *n* = 5 genotypes, D) Combination prevalences for a scenario with weak (*k* ≈ 0.02) and E) with strong interaction (*k* = ≈ − 0.41). In A, black dots show data from 5, 412 sexually active women in the Costa Rica Vaccine Trial reported by Chaturvedi et al. (2011) and lines show maximum likelihood fits performed using the `bbmle` package in `R` (Bolker, 2008). In B, the *β* and *γ* indicate transmission and recovery rates. In C, each circle indicates a prevalence (per genotype, per rank or per combination) that can be used as a summary statistics. In D and E, the shading indicates the infection rank (or number of coinfecting genotypes) and the class is a binary code indicating the genotypes present. We assume that genotypes B and E are the LR and A, C and D are the HR.

Fenton et al. (2014) compared several techniques using a dataset involving 2 species for which real within-host interactions were known from laboratory experiments. They concluded that correlation techniques performed worse and that the best method required time series and not just cross-sectional data (see (Shrestha et al., 2011) on how to infer interaction parameters from time series using particle filtering techniques). This is consistent with longitudinal data being generally richer for epidemiological inference than equilibrium data (Rohani & King, 2010). However, the restricted number of strain they used also potentially limited the power of their conclusion (3 ranks and 2 total prevalences versus 4 combinations).

### Parasite combination prevalences

Intuitively, there should be more information in the prevalence of each combination of genotypes than in the rank prevalence. With 5 circulating genotypes, there are only 6 possible ranks whereas there are 32 possible genotype combinations (Fig 1C). Earlier studies have already thought about using this data to compensate for the lack of longitudinal data. In particular, Vaumourin et al. (2014) considered systems with a larger number of genotypes using a variety of existing techniques (generalised chi-square, network models and multinomial GLM approaches) and developed a new association screening approach that has the advantage to identify and rank combinations based on their deviation from the expectation (see the Methods). Essentially, their methods consists in testing whether the observed genotype combination prevalence distribution significantly differs from the ‘neutral’ distribution in which parasites do not interact in their host (also referred to as ‘*H*_0_’). This neutral distribution is built from the total prevalence of each genotype assuming a multinomial distribution. As the Poisson distribution used by (Chaturvedi et al., 2011), it implicitly assumes an SI model with co-transmission.

One of the limitations of not having an explicit epidemiological model is that any type of heterogeneity into the system may lead to a deviation from *H*_0_. In particular, infected hosts may differ in their phenotypes for reasons other than the nature of the genotype(s) infecting them. Detecting an effect of interactions between genotypes on equilibrium prevalences therefore requires ruling out other important sources of host heterogeneity.

### Inference using explicit modelling

Our goal in this study is twofold. First, we want to assess the additional information that can be obtained from genotype combination data. Second, we also want to control for another source of host heterogeneity, namely the fact that some hosts may act as ‘super-spreaders’ (Lloyd-Smith et al., 2005). As mentioned above (Chaturvedi et al., 2011), these hosts should be more exposed to the infection and therefore have higher infection ranks independently of any features of the parasites themselves. Our hypothesis is that using a mathematical model that captures the epidemiological dynamics of *n* parasite genotypes (or species) in their 2^*n*^ coinfected host classes can allow us to address both our goals simultaneously.

Although our approach can be applied to many systems, we focus here on genital infections caused by different types of human papillomaviruses (HPVs) for several reasons. First, multiple infections between HPV types are common (Fig 1A) and well described thanks to screening for HPV-induced cancers (Vaccarella et al., 2010, Chaturvedi et al., 2011, Dickson et al., 2013). Second, their prevalences are relatively stable through time (Alemany et al., 2014). Third, HPV evolutionary rates are generally slow, which limits within-host evolution and facilitates detection (Bravo et al., 2010). Fourth, the existence of within-host interactions between HPV types is strongly debated, especially in the context of vaccination, given that they may affect a potential parasite evolutionary response (Murall et al., 2015).

Because of the high prevalence of coinfections and, more generally, because of the low immunogenicity and low pathogenesis of acute HPV infections (Alizon et al., 2017), many believe HPV between-types interactions in coinfected hosts to be negligible. However, pre-vaccine and vaccine studies have shown that there is limited natural cross-reactivity between phylogenetically related HPV types and that vaccines confer partial cross-immunity against non-target types (Herrero, 2009, Wheeler et al., 2012, Beachler et al., 2016). This means that there could be apparent competition mediated by the immune system. At the cellular level, recent data supports the existence of superinfection, that is one HPV type excluding the other from the cell (Biryukov & Meyers, 2018). For some types, virus loads also seem to differ in single and in coinfections (Xi et al., 2009), which could impact the host transmission and recovery rates. There is also indirect epidemiological evidence. First, infection by HPV is known to affect the risk of contracting another infection (Rousseau et al., 2001, Méndez et al., 2005, Tota et al., 2016) and to decrease the recovery rate of another type after coinfection (Trottier et al., 2008). Second, HPV coinfections may interfere with chronic infection and cancer. For example, when oncogenic ‘high-risk’ (HR) HPV types coinfect with non-oncogenic ‘low-risk’ (LR) types, time to diagnosis is longer and the risk of progression to cancer is lower (Sundström et al., 2015).

In summary, there are reasons to hypothesise that HPV types might interact when coinfecting a host and that these interactions could be large enough to affect the prevalence of some genotype combinations. Detecting or ruling out such interactions would also have a strong impact in the field. Importantly, our approach has no explicit within-host component and is therefore unable to detect a specific interaction. Instead, what it can detect is the overall effect of all the potential within-host interactions between genotypes.

As explained in the model section, it would be impossible to fit an interaction parameter between each HPV type. Instead, we sort HPV types into two groups and test for the existence of an interaction between HPVs belonging to these groups. Biologically speaking, the groups could correspond to HR and LR HPV types. Another possibility would be to compare HPV16 and HPV18, which together account for the vast majority of HPV-driven cancers, to the other HPV types.

To detect interactions between two groups of HPVs, we adopt mechanistic approach and simulate epidemiological coinfection dynamics. This is made possible by a recent analytical framework that can handle an arbitrary number of genotypes (Sofonea et al., 2015). In order to assess the ability to infer interactions from the observed coinfection classes, we use a regression-based Approximate Bayesian Computing (ABC) approach (Csilléry et al., 2012, Saulnier et al., 2017). We show that our method performs well on simulated data and can distinguish overall genotype interactions even in the presence of host behavioural heterogeneity.

## Methods

### The epidemiological model

The model is based on the deterministic ODE-based framework introduced by Sofonea et al. (2015) that allows for an arbitrary number of parasite genotypes to circulate in a host population without assuming any particular infection pattern (see Sofonea et al. (2017) for the importance of this relaxation). Furthermore, the framework enables cotransmission in the sense that infected hosts can simultaneously transmit any subset of genotypes they are infected with.

#### Multiple infections

Let us consider that hosts can be potentially infected by any combination of *n* parasite genotypes and sort them in classes according to the genotypes present (we use a binary code to map the presence/absence of the genotypes the hosts class labels). For computational reasons, we assume in the simulations that *n* ≤ 5, as the number of classes increases geometrically with the number of genotypes.

Epidemiological dynamics follow a classical susceptible-infected-susceptible (SIS) framework, where upon contact with an infected host, a ‘recipient’ host can acquire any subset of the genotypes carried by this ‘donor’ host (cotransmission). In terms of recovery, we assume that genotypes can be cleared independently. Importantly, each genotype *g* is cleared at a specific rate *γ*_*g*_ ≥ 1 year^−1^. This sets the average infection duration to a year (Insinga et al., 2007, Trottier et al., 2008). Given that we focus on HPV infections in young adults, we neglect infection-induced mortality.

Mathematically, the dynamics can be captured in a compact form using the master equation (Sofonea et al., 2015):

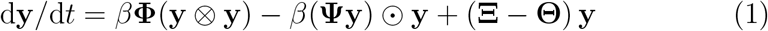

where **y** is the vector of densities of the 2^*n*^ host classes, ⊙ denotes the Hadamard (element-wise) matrix product, ⊗ the Kronecker (outer) product, **Φ**is the infection input flow matrix, **Ψ** is the infection output flow matrix, **Ξ** is the recovery input flow matrix and **Θ** is the recovery output flow matrix and *β* is the (constant) probability of transmission per contact that scales all infection processes. Equation system 1 allow us to track all the flows going in and out of host compartments through time. For simplicity, we neglect host demography (births and deaths) and assume that the host population size is constant. Given that infected hosts do not always sero-convert and that natural immunity is lower than vaccine-induced immunity (Beachler et al., 2016), we neglect immunisation in the model.

#### Population structure

The model was enhanced by splitting the host population into two sub-populations that differ in their contact rates (‘super-spreader’ versus ‘normal-spreader’ hosts) as shown in Figure 1B (Keeling & Rohani, 2008). Contacts between the two sub-populations follow a classical pattern based on the assortment (*a*) within host types, the proportion of each host type (*p*_1_ = *p* and *p*_2_ = 1 − *p*) and their activity rates (equal to *c*_1_ = 1 and *c*_2_ = *h*, with *h* ≥ 1). Overall, the contact rate between a ‘recipient’ individual from sub-population *j* and a ‘donor’ individual from sub-population *i* is

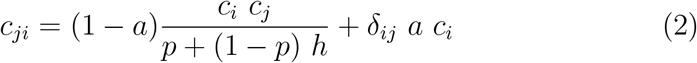

where *δ*_*ij*_ is the Kronecker delta and *h* is the difference in activity between the two host types.

This population structure implies that we have two vectors of host classes (**y**_**1**_ and **y**_**2**_). If we denote the combined vector **y**_•_ = (**y**_**1**_, **y**_**2**_), the master equation can be written similarly to 1 by updating the matrices in the following way:

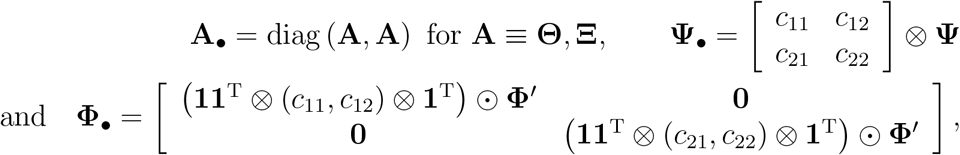

where **1** denotes the 2^*n*^-dimensional column vector with unit elements, and **Φ**^*′*^ is obtained byrepeating each 2^*n*^ × 2^*n*^ block **Φ**^[*i*]^ of the original 2^*n*^ × 2^2*n*^ matrix **Φ** = (**Φ**^[*i*]^)_*i*=1,…,2^*n*^_ as **Φ**^*′*^ = (**Φ**^[*i*]^, **Φ**^[*i*]^)_*i*=1,…,2^*n*^_.

#### Model simulations

The model was implemented and simulated in R. The script is already available upon request and will be published on a repository along with the part of the raw data (simulated prevalences).

The equilibrium prevalences from the deterministic model were used to generate datasets in finite populations of 1,000, 5,000 and 10,000 hosts assuming a multinomial distribution, where the probability to draw a host with a given genotype combination was equal to this combination’s prevalence.

#### HPV interactions

We neglected within-host dynamics and modelled the effect of genotype diversity on the infection parameters in the following way. First, we assumed that genotype transmission was unaffected by the presence of other genotypes in the host. This was motivated by the very high transmission probability of HPV per contact (Winer et al., 2006). Second, we assumed that interactions between HPV types take place through the recovery rates.

Even with 5 genotypes, this could mean 20 interaction parameters (e.g. how the presence of genotype *A* affect the clearance rate of genotype *B*). To reduce this complexity, we assumed that genotypes could be sorted into two groups (see the Introduction). Whenever a genotype from the second group coinfects a host with a genotype from the other group, its individual recovery rate is multiplied by a factor 1 + *k*, with *k* [0.5, 0.5]. We assumed that if there were several genotypes from the other group, the factor was still 1 + *k*. Genotypes from the first group were assumed to be unaffected by the presence of other genotypes (otherwise we would need an additional parameter and assumptions as to the interaction between the two parameters). If *k* is greater than 0, we expect host classes containing genotypes from the second group to be under-represented. The reverse is true if *k* is lower than 0. We assumed that one of the groups contained 3 genotypes and the other 2. We do not expect a different partitioning to affect the results and the exact partitioning should eventually be based on the data.

### Inference from distributions

In order to compare our framework to existing methods, we use 3 of the 4 techniques implemented by Vaumourin et al. (2014) in R. These are briefly described here but readers interested in more detailed should refer to the original publication. For each of these techniques, we analysed a dataset with two host types (normal-spreaders and super-spreaders) and a dataset with a unique host type. Our hypothesis is that these methods should not be able to distinguish between the heterogeneity caused by the genotype within-host interactions and that caused by host behaviour.

#### Association screening

This approach involves simulating datasets of occurrence count of each combination of gentoype based on the genotype prevalences (Vaumourin et al., 2014). From these simulations, a 95% confidence envelope is calculated for each combination, thus allowing to detect deviation from the expected distribution in the dataset (also referred to as *H*_0_).

#### Multinomial GLM

This model consists in calculating the deviance from a statistical distribution obtained with a Generalised Linear Model and a multinomial family. Practically, the multinomial logistic regression model was performed using the *vglm* function from the `VGAM` package in `R` (Yee, 2015).

#### Generalised chi-square

This test does not involve any simulations and is based on the expected chi-square distribution of the prevalence of each combination of genotype given the total prevalence of each genotype. Note that combinations found only in 5 hosts or less are grouped together.

### Regression-ABC

This method follows that developed in phylodynamics (Saulnier et al., 2017). In short, Approximate Bayesian Computation (ABC) is a likelihood-free method to infer parameter values from a given dataset (Beaumont, 2010). It consists in simulating many datasets, for which by definition the underlying parameters are known, and comparing them to the target dataset, the parameters of which we want to estimate. This comparison is often done by breaking the datasets into summary statistics. We use regression-ABC (Csilléry et al., 2012), which is divided into two steps. First, in the rejection step, only the simulated runs that are close enough from the target are kept. Second, a regression model is learnt on the remaining runs. Once we know how to map summary statistics to the parameter space, we can infer the parameters from any target dataset from which the same summary statistics can be extracts.

Using equation system (1) we calculated the equilibrium prevalences of each of the 64 host classes (32 classes for each host type) for 50,001 parameter sets. We used large and uninformative priors for the parameters (Figure S2). More specifically, we varied the competition intensity (our parameter of interest, *k* ∈ [−0.5, 0.5]) the transmission rate (*β* ∈ [0.5, 1.5]), the assortativity (*a* ∈ [0, 1]), the activity difference between host types (*h* ∈ [2, 20]) and the modifiers for the genotype-specific infection durations (*d*_*i*_ ∈ [0.6, 1], with the normalisation *d*_1_ = 1).

We compare three sets of summary statistics:

- the ranks set, which includes the 5 rank prevalences and the 5 total prevalence of each genotype, that is 10 summary statistics
- the comb set, which includes the rank set and the prevalences of the 32 combinations of genotypes, that is 42 summary statistics
- the all set, which includes the comb set for each of the two types of hosts (84 summary statistics) plus all the differences between each combination prevalence and its corresponding rank prevalence (64 summary statistics), that is 148 summary statistics.

The first set is intended to mimic an approach that would ignore combinations of genotypes (but that would capture host heterogeneity with super-spreaders). The second set is based on the type of data that could readily be accessed. The third is for a most optimistic scenario in which we would know which group each host belongs to. Importantly, we are using the same information used by earlier methods based on the prevalences of the genotype combinations. The only difference is that we combine some of these prevalences to generate additional summary statistics.

We compared several levels of tolerance using a preliminary run of the model (with narrower priors) and identified 50% as an optimal cut-off for the rejection: lowering the tolerance did not improve the inference (measured via the fraction of runs where the target value ended up in the 95% HPD), whereas increasing it decreased the inference quality.

Following an earlier study (Saulnier et al., 2017), we used a LASSO regression to learn the model. Although it performs a linear regression, it has the advantage to be less prone to over-fitting than more elaborate non-linear regressions, such as Support Vector Machines, neural networks or random forests. The LASSO adjustment was implemented using the glmnet R package and the ABC itself was performed using the abc package. In practice, one of the 50,001 runs was removed and used as a target, whereas the remaining runs were used to learn the regression model (after performing a rejection step). We repeated the operation 100 times to generate 100 target datasets. For completeness, we also analysed 100 runs with only a single host type to compare our method to existing ones and investigate the robustness of the ABC to a mismatch between the model used to simulate the target model and the one used to learn the regression model.

## Results

### Associations and competition intensity

We hypothesised that current methods, which implicitly assume a simple SI epidemiological model with cotransmission, may have difficulties to detect within-host competition between HPVs if there is another source of host heterogeneity than coinfection status. To test this hypothesis, we used our model to simulate target sets of genotype combination prevalences for known parameter values.

Figure 2 shows the performance of the association screening approach conceived by Vaumourin et al. (2014). With two host types, ‘normal-spreaders’ and ‘super-spreaders’, the number of significant interactions, i.e. the number of host types that show a prevalence that departs from the neutral expectation (*H*_0_), is independent from the intensity of the competitive interactions, |*k*| (Fig. 2A). Furthermore, the fraction of these predictions that correspond to what the analytical model would predict based on the nature of the interaction, i.e. the sign *k*, is always close to 50% (Fig. 2C). On the contrary, if we assume that there are no super-spreaders, then the number of significant interactions increases with competition intensity (Fig. 2B). The proportion of correct predictions also increases with competition intensity to reach a maximum estimated median of above 75% (Fig. 2D). This suggests that this method can be appropriate to detect strong competitive interactions in homogeneous host populations.

**Fig 2.**
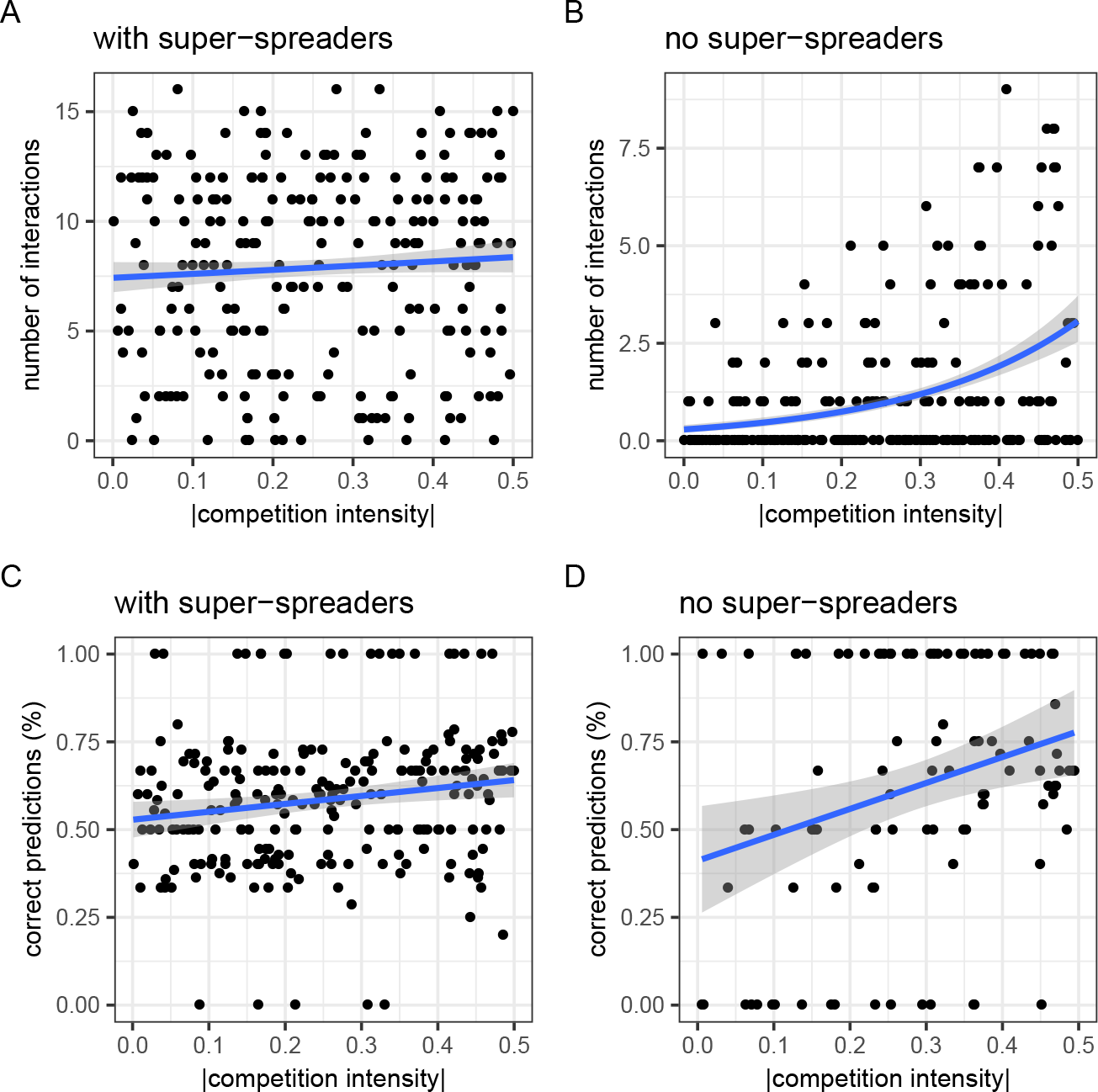
Total number of interactions detected with the association screening method (A and B) and fraction of these interactions that are consistent with model predictions (C and D). This analysis is ran for a model with two host types (A and C) or a single host type (B and D). The blue lines show the result of a linear model fit (A and B) and generalised linear model fit assuming a Poisson distribution of the outcome variable (C and D). Grey areas are prediction intervals based on the standard error of the fit. In panels A and C, *h* = 1 and *a* = 0. We assume that there are *N* = 5, 000 hosts in the population.

The Chi-square and GLM approaches are more qualitative: they either detect a difference with *H*_0_ or not. In Supplementary Figure S8, we show that the GLM fails in both cases. For the chi-square approach, we do detect an increasing probability that the test is significant with increasing competition intensities (|*k*|) with a maximum of ≈ 70%. As we will see later on, analysing the same target datasets with the ABC approach yields very different patterns.

### Epidemiological model: single runs

We first show the prevalences of combination of genotypes in two scenarios, one with moderate interactions (parameter set #2 with the competition intensity parameter *k* ≈ 0.02, Fig. 1D) and another with strong interactions (parameter set #7 with *k* ≈ −0.41, Fig. 1E). When the interactions are weak, we clearly see the different ranks: uninfected hosts are on the top, then there is a row with the five singly infected host types, etc. When competition intensity increases, these ranks become impossible to distinguish. Figure 1D also illustrates that each parasite genotype in this model has its own infection duration, since they do not all have the same prevalence in single infection (see rank 1 point data). Importantly, we only show the total prevalence of each combination but these may differ among each of the two host types (in the ‘super-spreader’ population high rank genotype combinations are more prevalent).

Our goal is to infer the intensity and sign of the interaction between HR and LR genotypes (parameter *k*) in such a heterogeneous host population. To this end, we applied an ABC approach. As any bayesian method, this means searching a prior distribution in the parameter space. This distribution is shown for all the key parameters in Figure S2. We drew 50,001 parameter sets in this prior, used them to simulate equilibrium densities similar to the ones shown in Figures 1D and E.

Figure 3 shows the results for parameter set #3 and illustrates how using more summary statistics helps to narrow the distribution from the prior for a dataset with 10,000 individuals. If we only use the ranks, we do narrow the prior distribution but its width remains large enough such that 0 (no interaction) cannot be ruled out from the 95% Highest Posterior Density (HPD), which can be seen as a credibility interval (Fig. 3B). Using the prevalence of the genotype combinations in addition to the prevalence of the infection ranks as summary statistics for the ABC allows us to narrow this interval and to exclude 0 from the 95% confidence interval (Fig. 3C). Using additional information, for example being able to distinguish between the two host types, would narrow it even more as we will see below.

**Fig 3.**
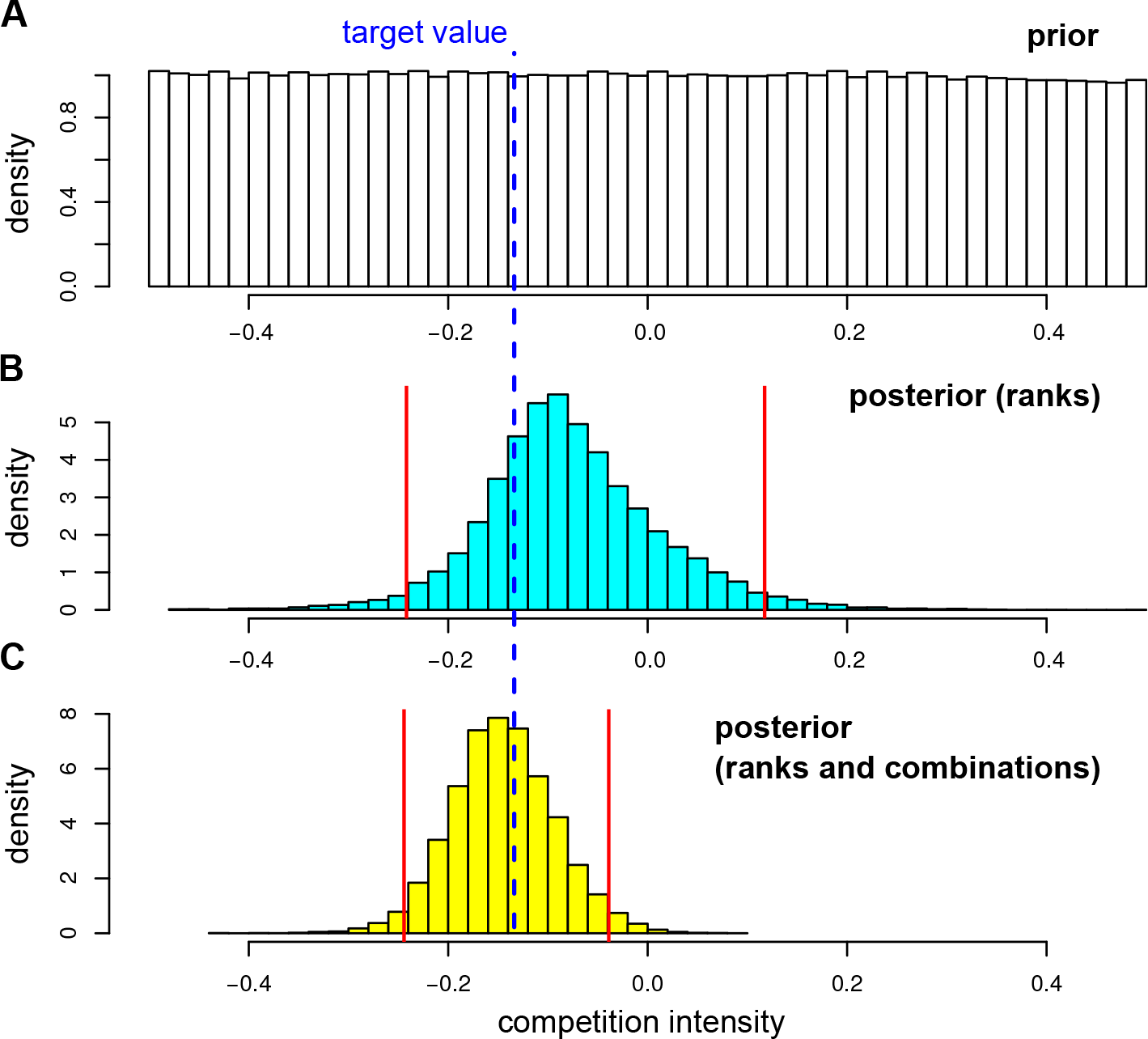
Inferring competition intensity (*k*). Prior (A) and posterior distributions using the ranks (B) or the comb set (C) of summary statistics. The dashed blue line shows the target value (*k* ≈ −0.13) and the red lines the 95% Highest Posterior Density (HPD).

### Epidemiological model: cross-validation

The previous analysis was based on a single set of target parameters. Since all parameters may vary in a relatively large prior distribution (Fig S2) and since *k* may be easier to infer in some settings, we assessed the performance of the ABC approach following a leave-one-out cross-validation procedure, where we treated one simulation as observed data and the remaining as learning data. We varied the number of sampled individuals and used 100 targets for each. Furthermore, we analyse a third set of summary statistics involving the prevalences of infection ranks and genotype combinations for the two hosts sub-populations (see the Methods).

As expected, the width of the 95% HPD for the estimate of competition intensity decreased with the number of host sampled (Fig. 4A). On the same figure, we see that including more summary statistics also decreased the width of this interval, especially for an infinite sample size.

**Fig 4.**
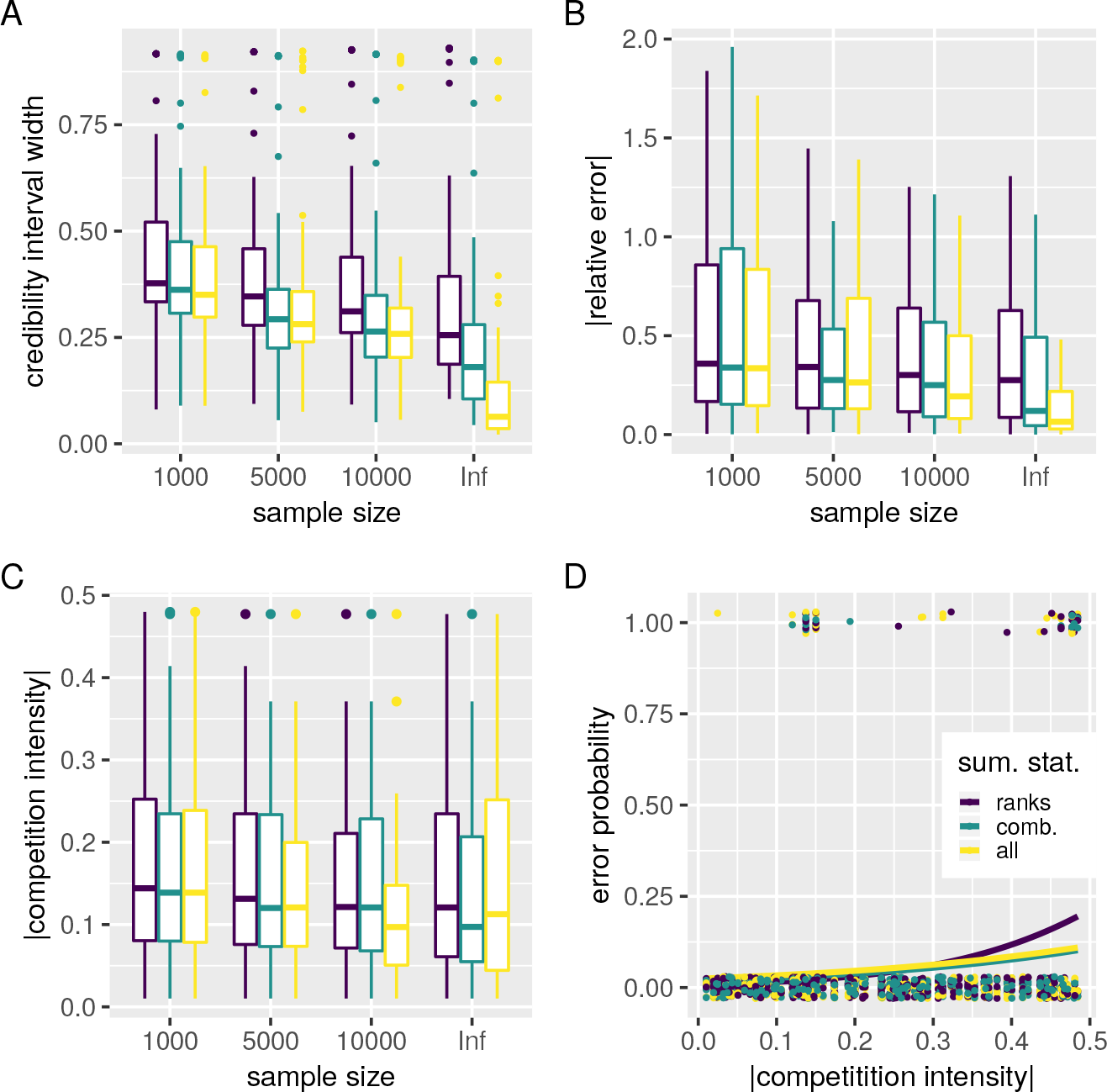
ABC inference precision over 100 runs. A) 95% Highest Posterior Density (HPD), B) absolute value of the relative error, C) average of the absolute value of competition intensity in runs where 0 is in the 95% HPD and D) runs for which the target value lies outside the 95% HPD. Colours indicate the summary statistics used for the ABC. In D, the lines show the result generalised linear models fits assuming a binomial distribution of the outcome variable.

In terms of the relative error made when estimating the competition intensity parameter (*k*), we found a similar effect with a lower error when more hosts were sampled or more summary statistics were involved (Fig 4B). This effect is the clearest when using all the summary statistics in an infinite population. In general, we see that increasing the number of summary statistics does not help when few hosts are sampled (all three sets are similar when *N* = 1, 000) and that using the prevalences of the genotype combinations only improves the analysis if enough hosts are sampled (5,000 or more). In general, the relative error decreased with competition intensity (figure not shown).

If we focus on the runs for which we could not exclude an absence of interaction (i.e. *k* = 0 lied within the 95% HPD), we see that the number of such runs decreased as the number of summary statistics increased (Fig S6). We also see that, in these runs, competition intensity decreased with the sample size and with the number of summary statistics involved (Fig. 4C). Notice that for large sample sizes, 95% HPD are narrower, which makes it more difficult to exclude an absence of competitive interactions.

Finally, the probability to make an error in the inference, which we define as having the target value outside the 95% HPD, was close to the expected 5% (6.25% with the ranks and 5% with the comb sets). This probability slightly increased with competition intensity, especially when the genotype combination prevalences were ignored in the ABC (Fig. 4D). Therefore, we have the somehow unexpected result that genotype combination data is even more important to anayse datasets where competitive interactions are particularly strong.

### Removing host heterogeneity

We then used the ABC approach to reanalyse the target sets with a single host type shown in Figure 2B. This allowed us to do more than simply compare methods. Indeed, in our prior for the ABC, the heterogeneity parameter is greater than 2. This means there is a mismatch between the model we assumed for the ABC (2 host types with some heterogeneity between them) and that used to generate the target data (1 host type). We can therefore evaluate the robustness of the inference method to a small error in model specification.

We investigated the relationship between genotype competition intensity (*k*) and our ability to reject an absence of interaction (*k* = 0) from the 95% HPD in a situation with two host types and one host type in the target dataset. Priors were identical to the other analyses and shown in Figure S2. In both situations, cases where the true competition value was not in the 95% HPD interval were close to 5% as in the other runs. We then investigated how often an absence of competition (that is *k* = 0) could be rejected. This is similar to the *H*_0_ tested by Vaumourin et al. (2014). We found that we could detect competition for 55% of the target values in a model with super-spreaders and for 63% of the target values in model with only a single host type. In the latter we also made one error, i.e. inferred a positive interaction for a negative target. This is because in this specific parameter set, the modifiers for the infection duration of the two LR genotypes (*d*_2_ and *d*_5_) were low, whereas that of the HR were all high, therefore perfectly mimicking a competition interaction. Figure 5 also shows that, as expected, the ability to reject *H*_0_ increased with competition intensity. Overall, removing the heterogeneity in the data due to differences in host behaviour does increase our ability to detect competitive interactions.

**Fig 5.**
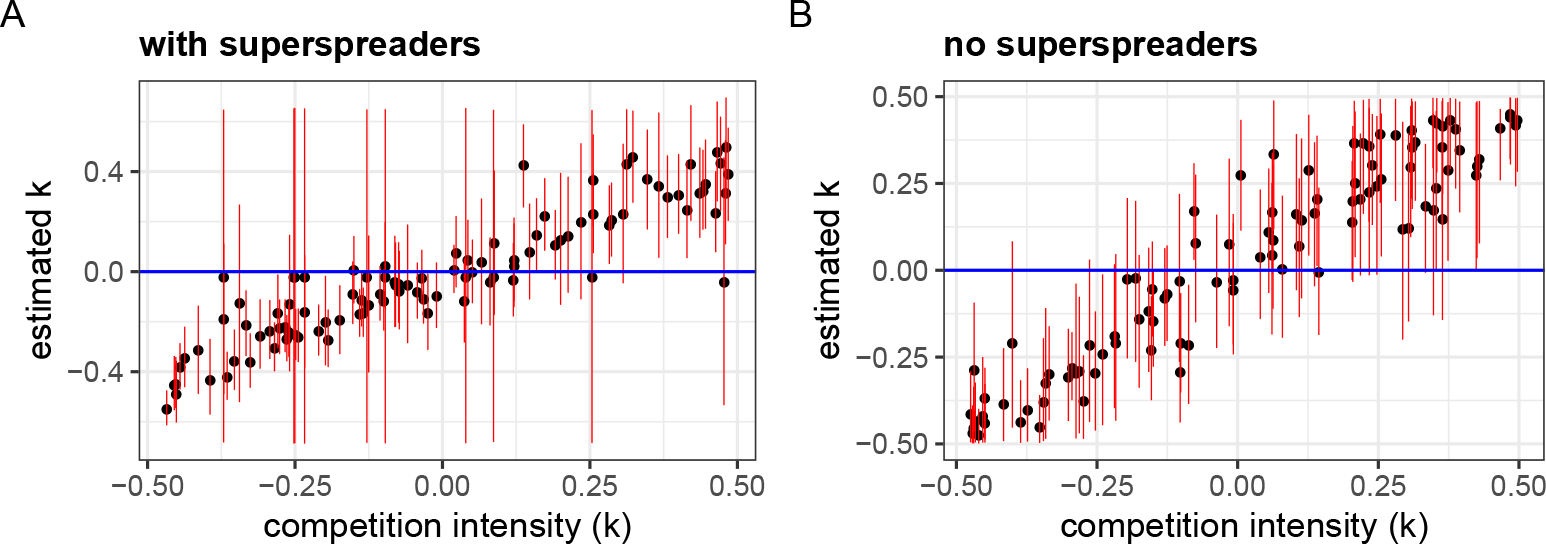
Inferring competition parameter (*k*) in, a setting with (A) and without (B) host behavioural heterogeneity. Red lines show the 95% credibility interval and the blue line shows the absence of interaction (*k* = 0). The target runs are identical to that in Figures 4 and 2 with *N* = 5, 000 hosts and the comb set of summary statistics.

## Discussion

Multiple infections are known to affect the virulence of an infection (Balmer & Tanner, 2011), the spread of infectious diseases (Abu-Raddad et al., 2006) and their evolution (Alizon et al., 2013). This is due to the fact that when sharing a host, parasites can interact in various ways such as competing for host resources, exploiting molecules they produce or even indirectly via cross-reactive immune response (Mideo, 2009). The goal of this study was to determine to what extent the prevalence of specific genotype combinations can inform us on the net effect of all these interactions.

By generating prevalence data from a mechanistic epidemiological model, we were able to first test the power of existing heuristic methods based on neutral distributions that implicitly assume a Susceptible-Infected (SI) model with co-transmission and only a single type of hosts. We showed that introducing host heterogeneity into the model can modify the distribution of genotype combination prevalences in a way that makes within-host interactions between genotypes largely undetectable. This therefore corroborates a limitation often mentioned in such studies, which is that departures from ‘neutral’ distributions (*H*_0_) need not be due to interaction between parasite genotypes.

We then used an ABC approach to infer parameters from the model. We show that this yields more consistent results than existing methods. As expected, the accuracy of the method increases with the number of hosts sampled. We also showed that using the prevalence of all the combinations of host classes tends to decrease the error made compared to using only the prevalence of infection ranks. Finally, adding information in the target data about host type (‘super-spreader’ or ‘normal-spreader’) can further improve the power of the inference.

The fact that decent results can be obtained by only using the rank of the infections may seem surprising considering the difficulty from existing models to infer interactions. This could mean that accounting for host behavioural heterogeneity is more important than adding additional information via the genotype combinations. Another reason could be that we here use the same model to generate the target dataset and the learning datasets, which facilitates the ABC inference. However, we do show that our inference method performs very well to infer competitive interactions when there is a slight mismatch between the true model and that used in the ABC.

As illustrated by Fig S7, the accuracy of the inference varied widely across parameters. For the interaction parameter (*k*), the inference reduced the initial 95% HPD of the prior by 66%. In comparison, this was less than for the transmission probability (*β*, 75%), but much better than for the assortativity parameter (*a*, 45%), host heterogeneity (*h*, 38%) or the individual recovery rates of each genotype *i* (*γ*_*i*_, 13%).

There are several ways to extend this framework. One would be to use more powerful non-linear machine learning regression techniques, such as neural networks. However, these may be more difficult to parameterise than the linear one we used. Furthermore, even though it contains several parameters, our model remains relatively simple compared to the power of these algorithms.

Here, we have also generally assumed that the epidemiological model is known. There are two ways to extend this. One can be to perform rigorous model comparison to see whether a simpler model (for instance with a single host type), might not fit the data better. This could be done readily using regression-ABC, for instance with random forests (Pudlo et al., 2016). Another extension would be to use an agent-based model with sophisticated agent behaviours to generate a richer dataset. This would be useful in itself to generate test runs with known parameter values to further test the power of our method on more noisy data. It would also allow to control for biases related to the contact network structure between hosts and the dynamical aspect of sexual partnerships that have been shown to interfere with the detection of coinfection interactions (Malagón et al., 2016).

Finally, the next step is, of course, to test this model using actual epidemiological data. Even in the case of HPV, analysing real data will require to add several processes we chose to ignore here. First, HPV detection tests may exhibit cross-reactivity between HPV types, thus inflating the prevalence of some genotype combinations. This effect if well described and can be handled for each detection test. Second, when hosts are infected by many HPV types, some of these may not be detected, thus decreasing the prevalence of high-rank infections. This effect is more subtle and would require to be inferred in the model.

Importantly, we focused here on HPV but other systems could be studied, in particular coinfections between different parasite species. However, it is important to stress that the underlying epidemiological model must be consistent with the the life-history of the parasite(s).

Overall, ABC and machine learning allow us to extract the information from the equilibrium prevalence of all the combinations of genotype prevalences. Therefore, combining coinfection modelling with epidemiological data can bring new elements to the controversy regarding the importance of interactions between HPV types.

## Supporting information

In addition to the supplementary figures, the archive SupplementaryMaterials.zip contains all the scripts used to generate the cross-validation results and plot the figure, along with a master table containing the results for each scenario (summary statistics used, tolerance, parameter ranges).

## Acknowledgments

We thank Elise Vaumourin for sharing her R script and helping with its implementation. We also thank Dustin Brisson, Erick Gagne and an anonymous reviewer from Peer Community in Ecology for helpful comments.

## A Supplementary Figures

**Fig S1.**
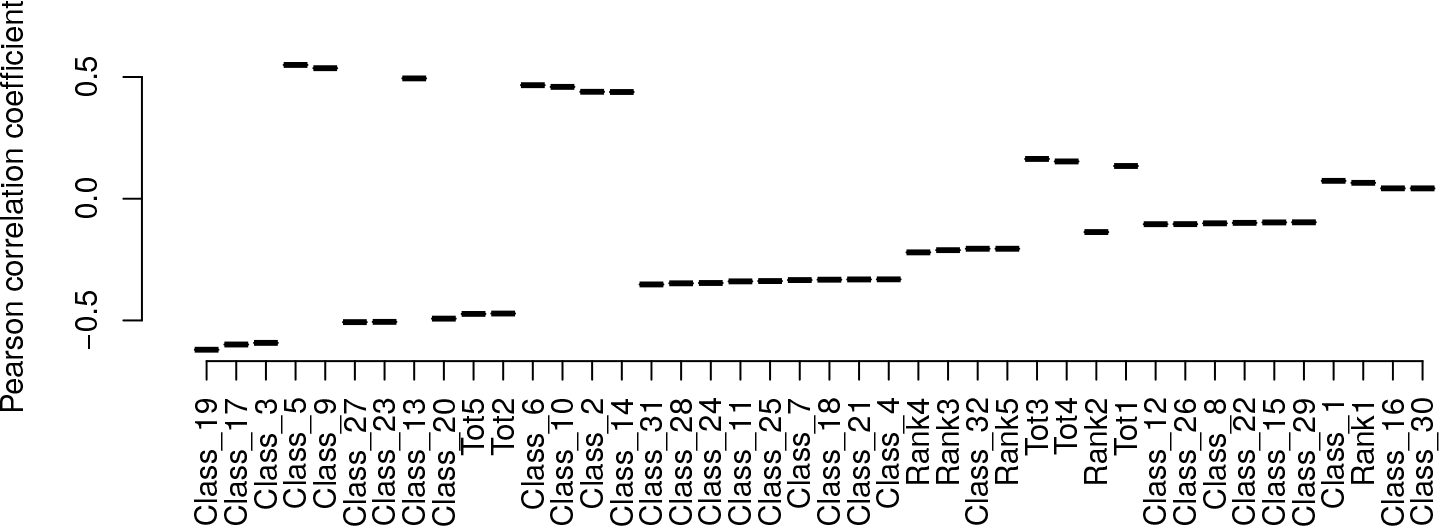
Correlation between competition intensity and combination, rank or genotype prevalence. The values show the Pearson correlation coefficient obtained using 1,000 parameter sets from the ABC training dataset (priors in Figure S2).

**Fig S2.**
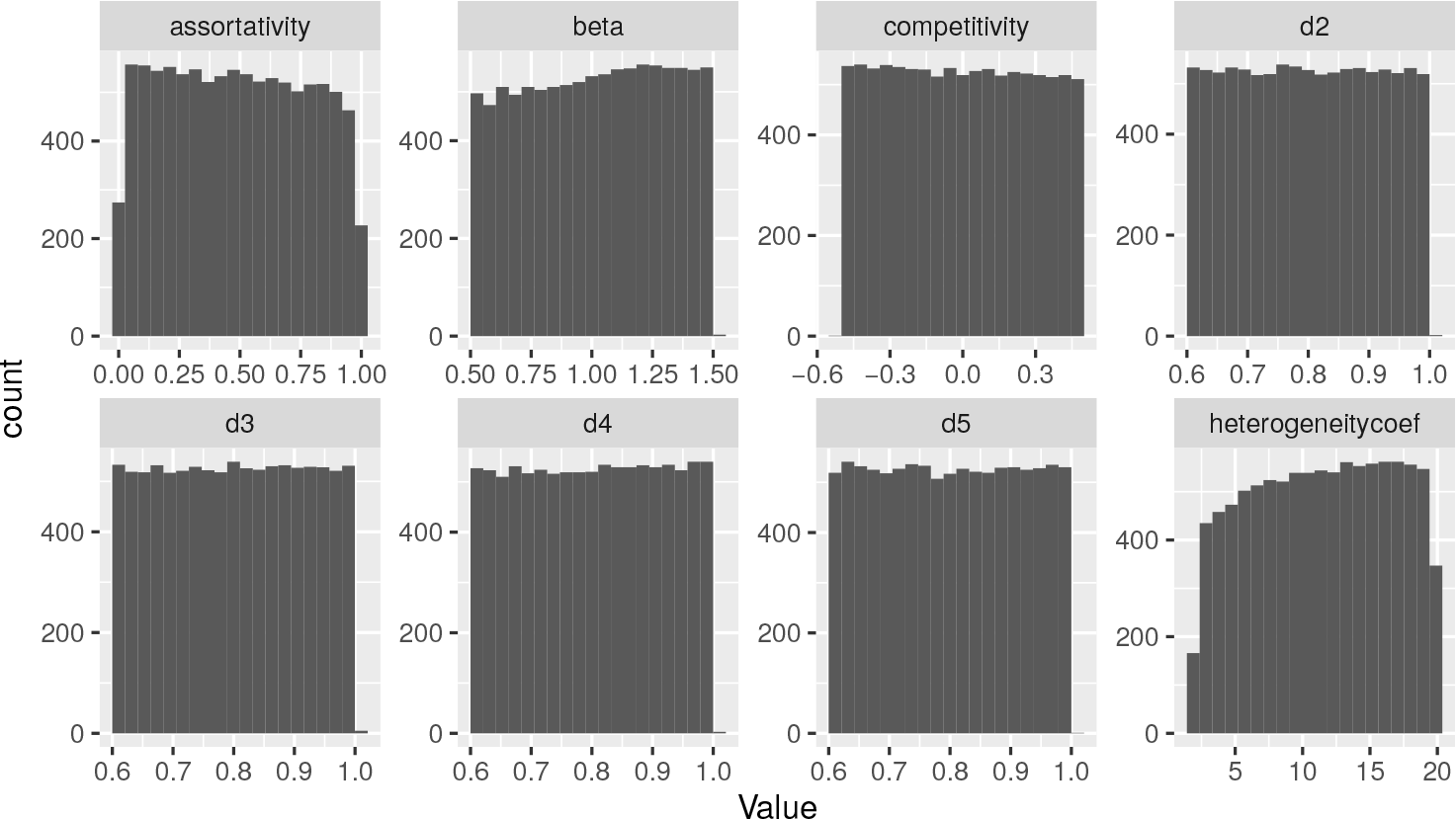
Prior distributions for all the parameters. The same priors are used to generate target datasets and training datasets.

**Fig S3.**
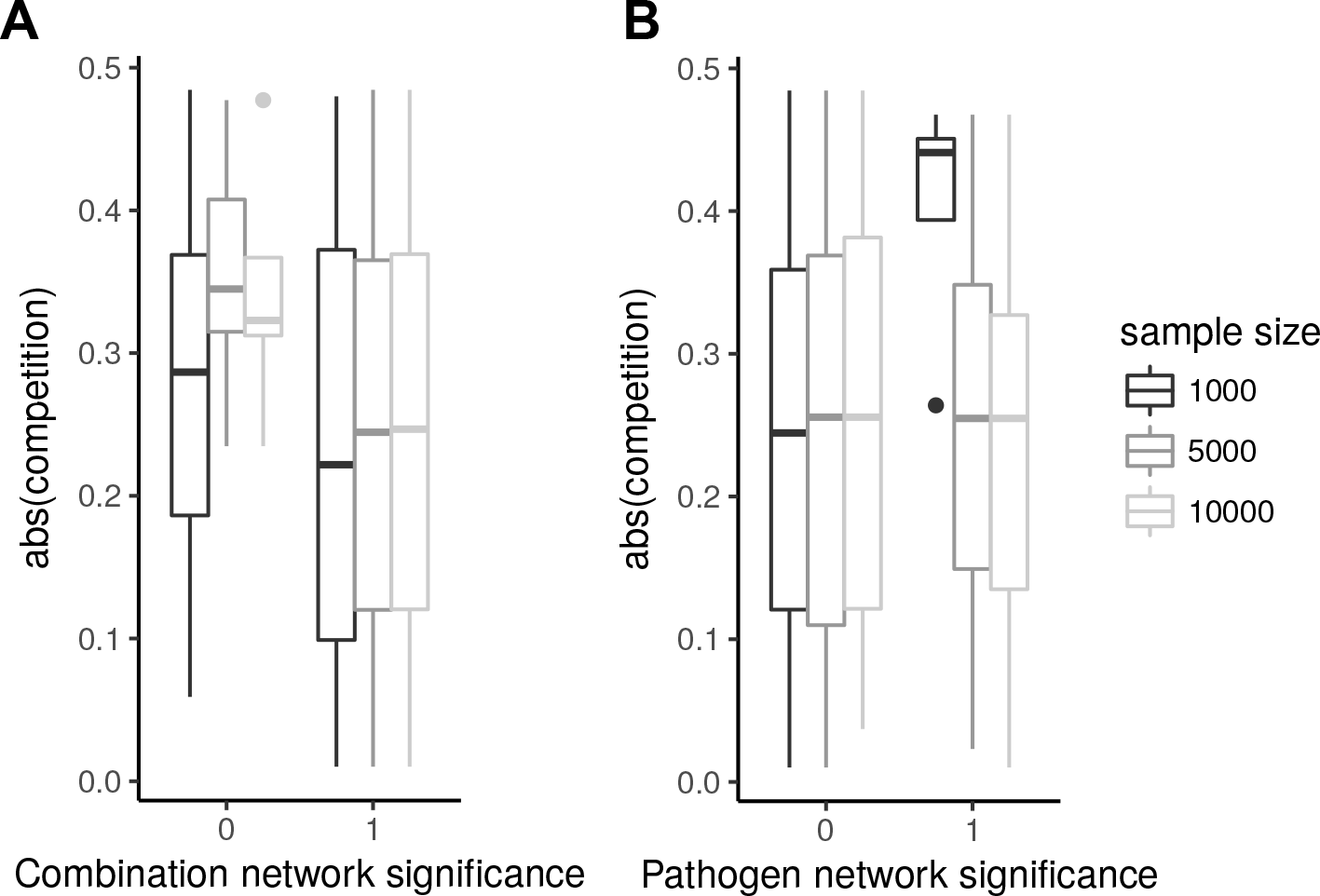
Difference in competition intensity depending on the p-value of the network-based test. A) If the combination network test is non significant, the interaction is likely to be strong. B) The difference for the pathogen network in the small sample size scenario is explained by the rarity of significant tests.

**Fig S4.**
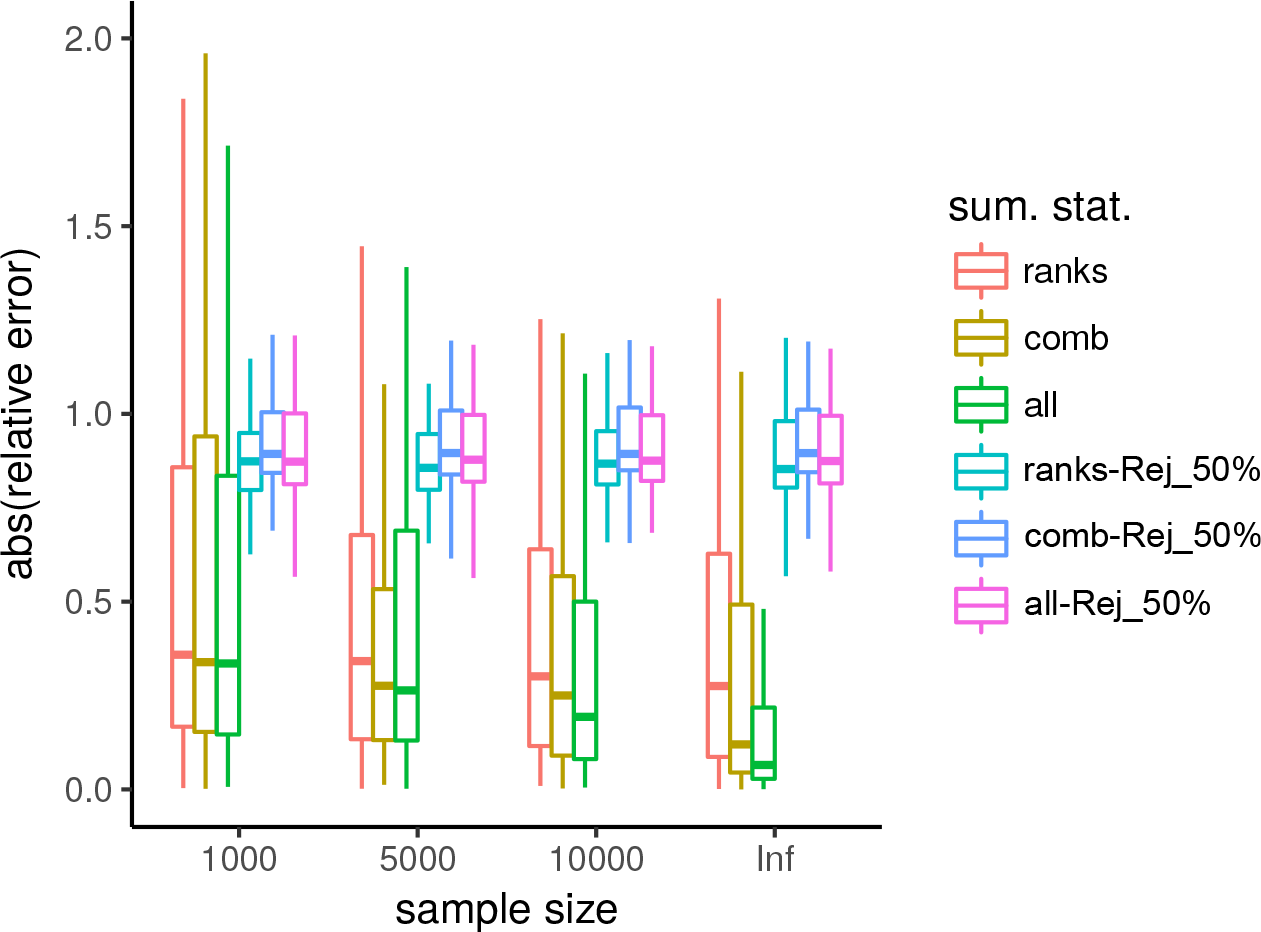
Relative error depending on the summary statistics and the methods used. The regression part of the ABC improves the inference compared to the rejection step alone.

**Fig S5.**
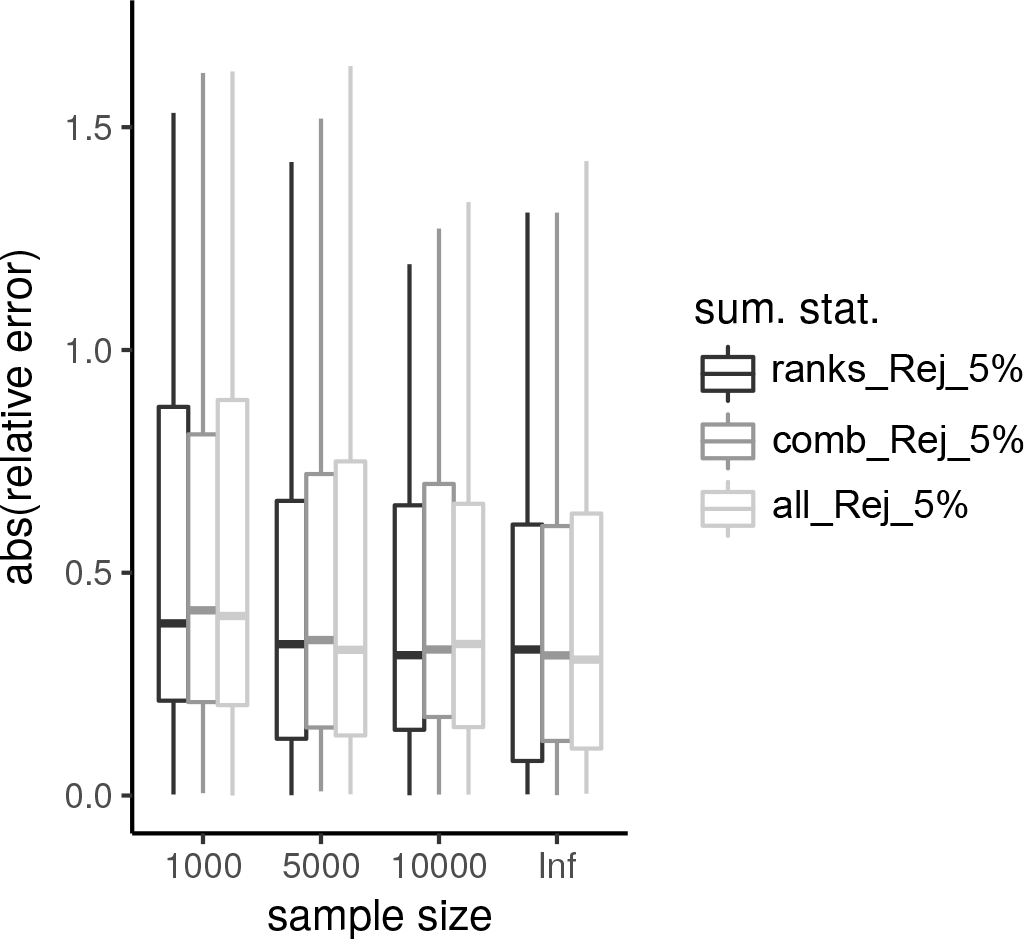
Rejection-based inference. When ignoring the regression part of the ABC, the set of summary statistics has little effect on the quality of the fit.

**Fig S6.**
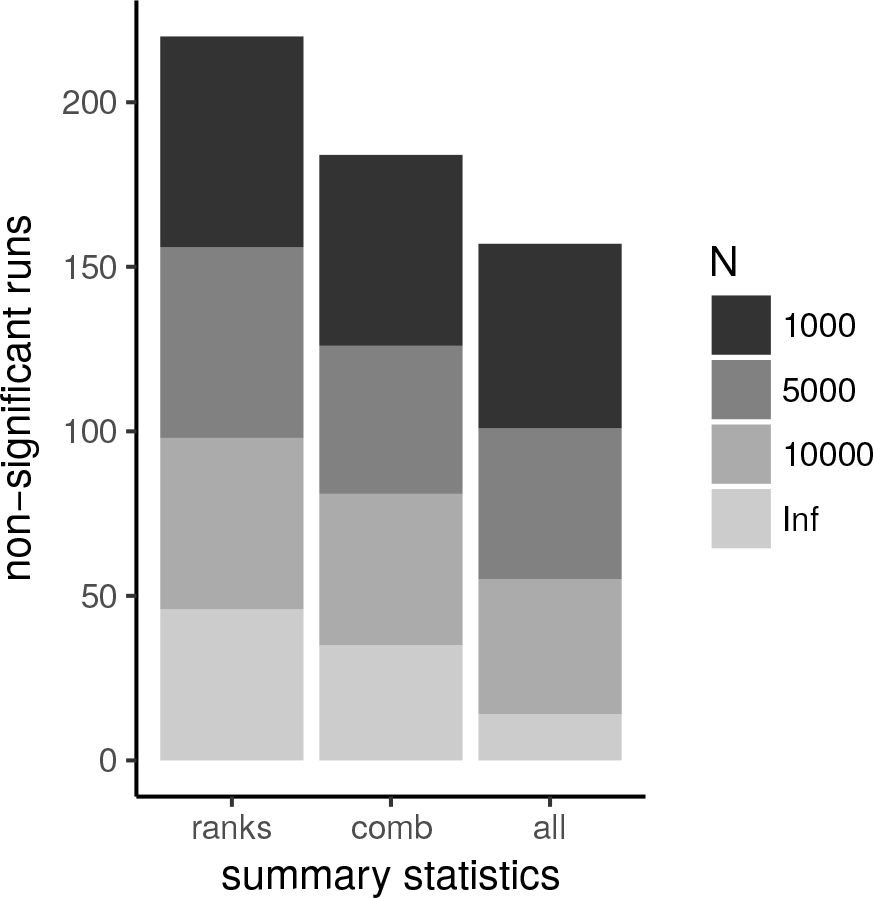
Number of runs where 0 cannot be excluded from the 95% HPD. Increasing the sample size and the number of summary statistics decreases the number of such non-significant runs.

**Fig S7.**
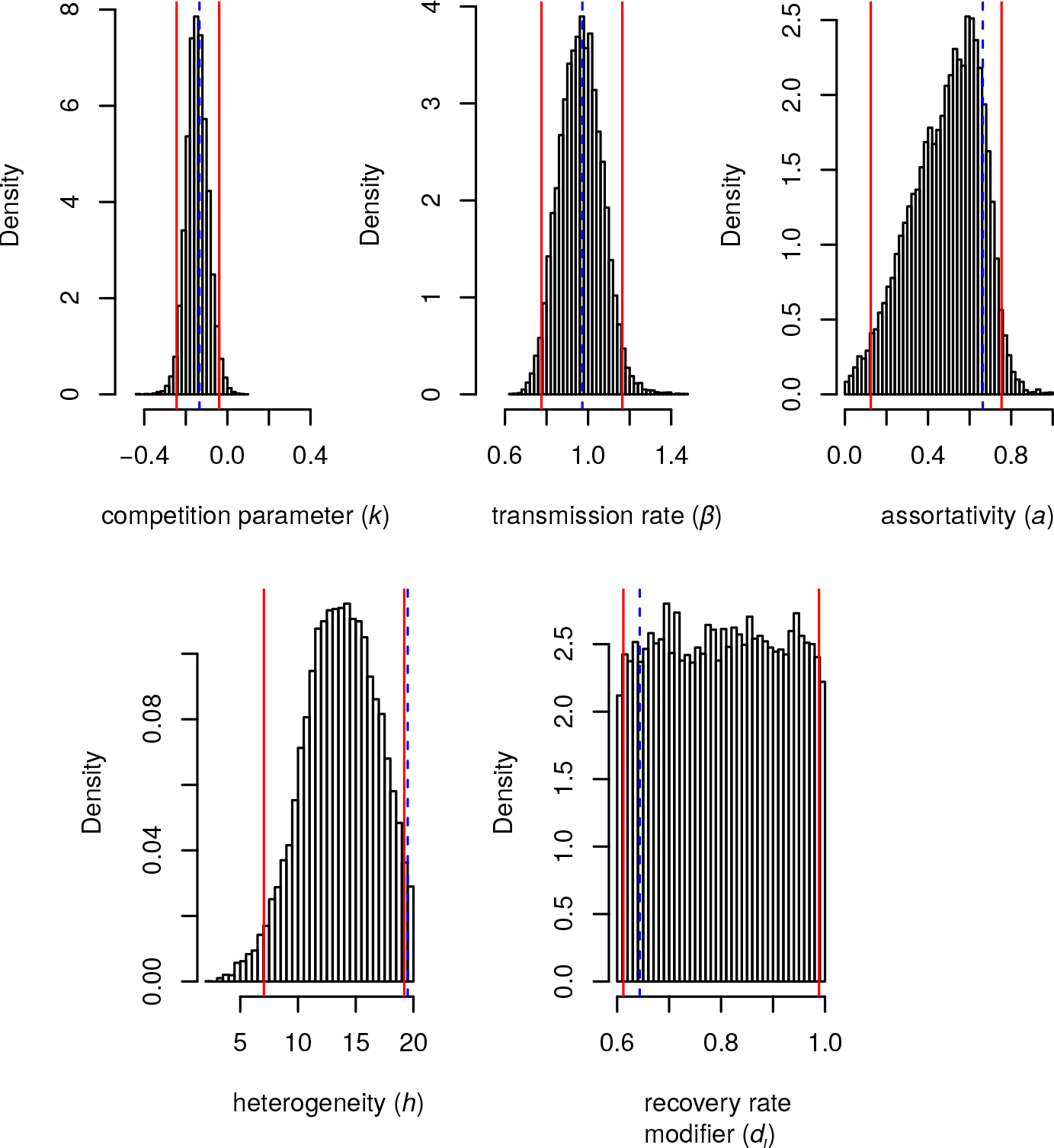
Inferring model parameters using the ranks and the combinations as summary statistics. We use parameter set #3 as the target and the remaining 50,000 sets to perform the ABC. The dashed blue lines show the target values and the red lines show the 95% Highest Posterior Density (HPD).

**Fig S8.**
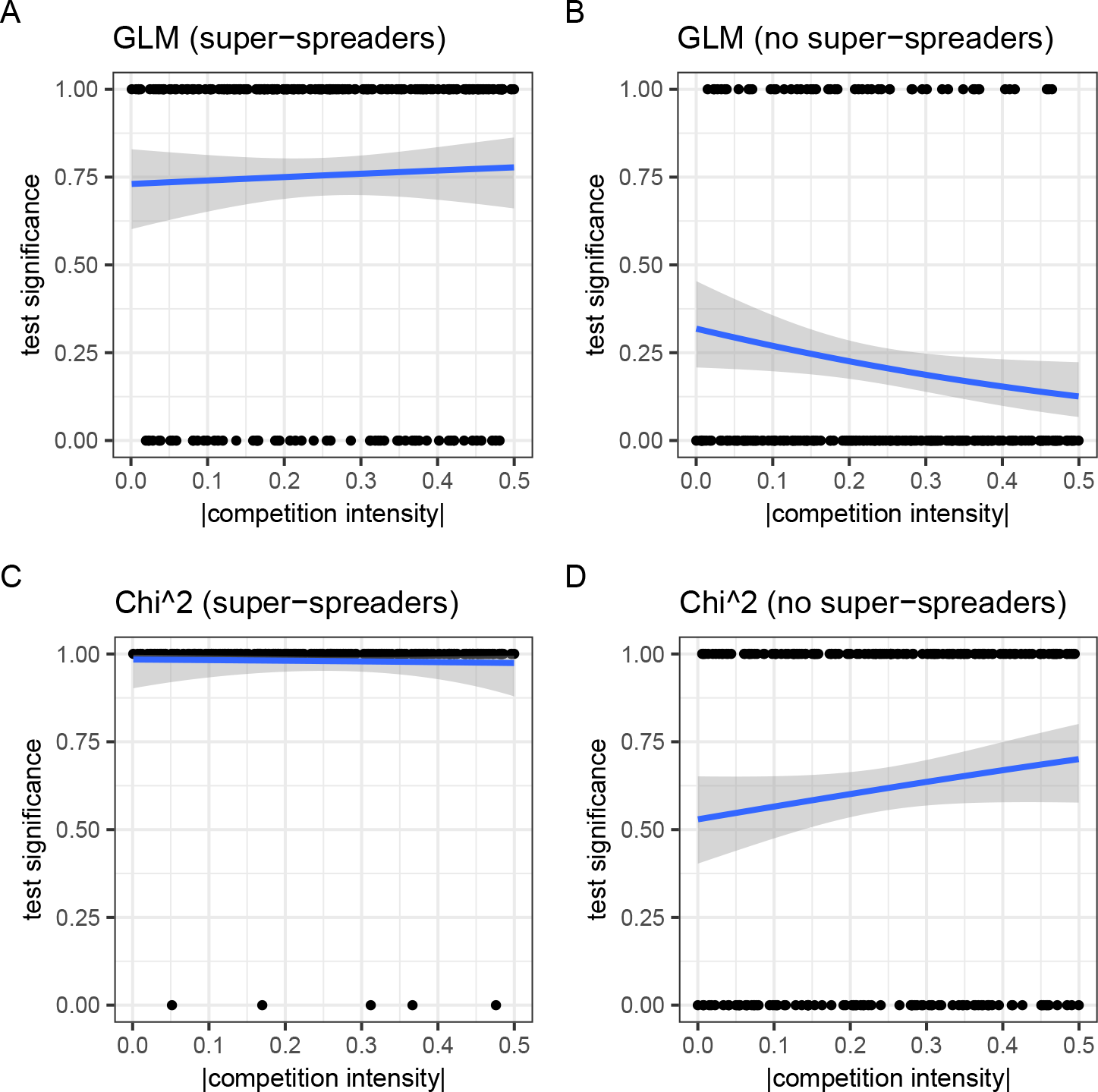
Significancy of the GLM (A and B) and the chi-square (C and D) approaches. This analysis is ran for a model with two host types (A and C) or a single host type (B and D). In panels A and C, *h* = 1 and *a* = 0.

